# Defective satellite DNA clustering into chromocenters underlies hybrid incompatibility in *Drosophila*

**DOI:** 10.1101/2021.04.16.440167

**Authors:** Madhav Jagannathan, Yukiko M. Yamashita

**Affiliations:** Department of Biology, ETH Zürich; Howard Hughes Medical Institute; Whitehead Institute and MIT

## Abstract

Although rapid evolution of pericentromeric satellite DNA repeats is theorized to promote hybrid incompatibility (HI)(*1*–*4*), how divergent repeats affect hybrid cells remains poorly understood. Recently, we demonstrated that sequence-specific DNA-binding proteins cluster satellite DNA from multiple chromosomes into ‘chromocenters’, thereby bundling chromosomes to maintain the entire genome in a single nucleus(*5, 6*). Here we show that ineffective clustering of divergent satellite DNA in the cells of *Drosophila* hybrids results in chromocenter disruption, associated micronuclei formation and tissue atrophy. We further demonstrate that previously identified HI factors trigger chromocenter disruption and micronuclei in hybrids, linking their function to a conserved cellular process. Together, we propose a unifying framework that explains how the widely observed satellite DNA divergence between closely related species can cause reproductive isolation.

## Main Text

### *Drosophila* hybrids exhibit chromocenter disruption and micronuclei formation

Our previous work showed the importance of pericentromeric satellite DNA in encapsulating the full complement of chromosomes into a single nucleus(*5, 6*). Sequence-specific DNA-binding proteins cluster their cognate pericentromeric satellite DNA repeats to create physical links between heterologous chromosomes, forming a cytological structure known as chromocenter (Fig. 1A). Depletion of satellite DNA-binding proteins led to the detachment of heterologous chromosomes from one another (chromocenter disruption), their subsequent loss from the primary nucleus (micronuclei) and cell death (Fig. 1B). Together, we demonstrated how identical satellite DNA repeats on multiple chromosomes are important for chromocenter formation and maintenance of the entire genome in the nucleus. This prompted us to hypothesize that the highly divergent satellite DNA repeats between closely related species may fail to form chromocenters properly, ultimately resulting in hybrid incompatibility (Fig. 1C). *Drosophila melanogaster* diverged ∼2-3 mya from the *Drosophila simulans* species complex (e.g. *D. simulans* and *D. mauritiana*)(*7*) but these species still maintain near complete synteny(*8, 9*). However, the satellite DNA content of *D. melanogaster* is markedly different from that of the *D. simulans* species complex (Fig. 1D) (*10, 11*). Most strikingly, the (AATAACATAG)_n_ satellite DNA whose clustering into chromocenters in *D. melanogaster* is required for viability(*6*), is completely absent in the *D. simulans* species complex (Fig. 1D). Rather, these species contain an unrelated repeat (GAACAGAACATGTTC)_n_ at the corresponding autosomal locations (Fig. 1D) (*10, 11*). In addition, the (AATAT)_n_ satellite DNA repeat, whose clustering is important for *D. melanogaster* fertility, exhibits differential abundance and chromosomal locations between these species (Fig. 1D) (*10, 11*).

**Figure 1.**
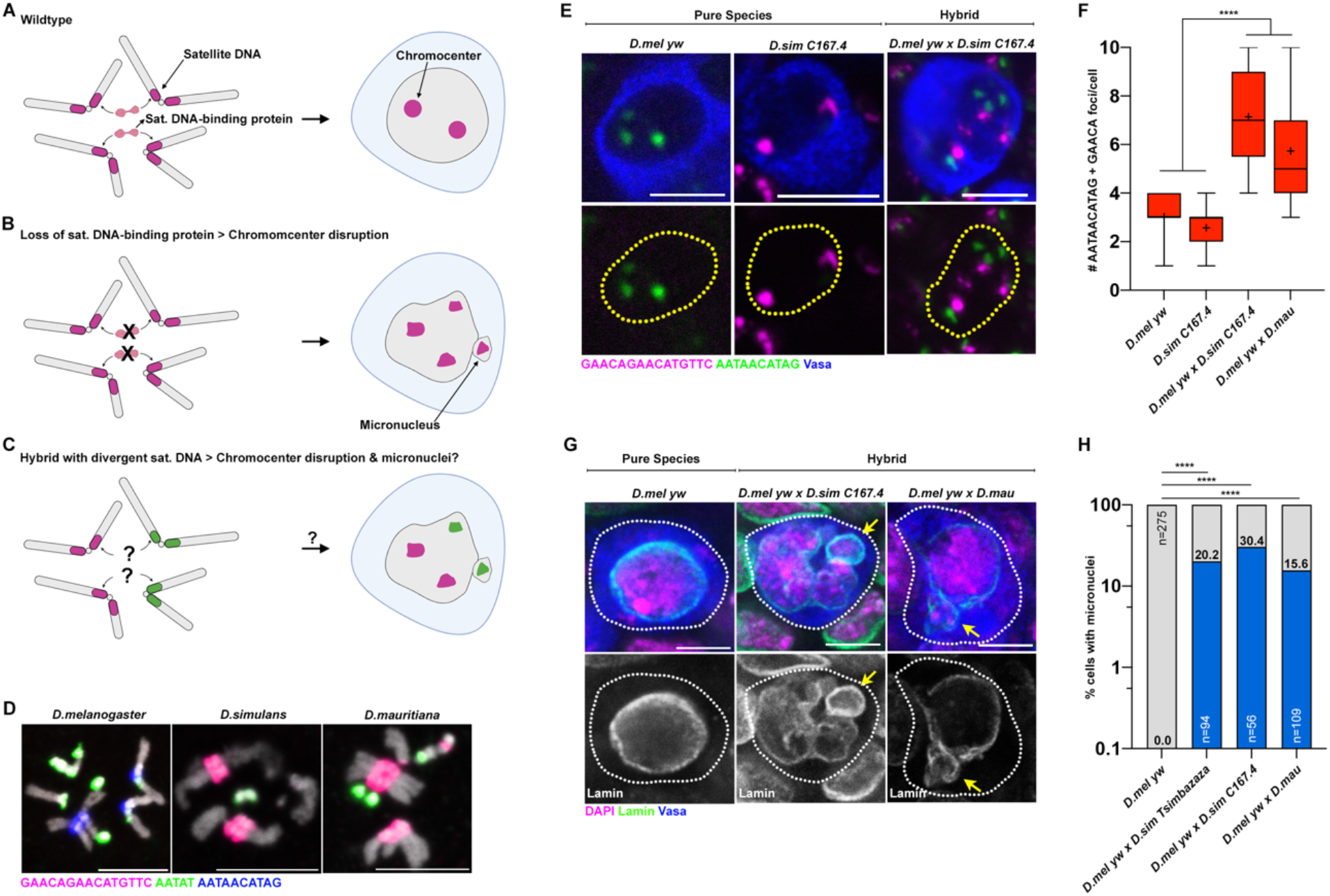
Chromocenter disruption and micronuclei in the larval germ cells of *D*.*melanogaster-D*.*simulans/D*.*mauritiana* hybrids. (A – C) A model of chromocenter formation (A) and function (B) in pure species and proposed dysfunction (C) in hybrids. (D) FISH against the (AATAACATAG)_n_ satellite (blue), the (AATAT)_n_ satellite (green) and the (GAACAGAACATGTTC)_n_ satellite (magenta) on larval neuroblast mitotic chromosomes from the indicated species and co-stained with DAPI (gray) (E) FISH against the (AATAACATAG)_n_ satellite (green) and the (GAACAGAACATGTTC)_n_ satellite (magenta) in the larval female germ cells from the indicated species and hybrids and co-stained with Vasa (blue) (F) Box-and-whisker plot of the total number of (AATAACATAG)_n_ and (GAACAGAACATGTTC)_n_ chromocenter foci per larval germ cell from *D*.*mel yw* females (n=38), *D*.*sim C167*.*4* females (n=28), *D*.*mel yw* x *C167*.*4* hybrid females (n=21) and *D*.*mel yw* x *mau* hybrid females (n=42). **** represents p<0.0001 based on Tukey’s multiple comparisons test from an ordinary one-way ANOVA and crosshairs mark the mean (G) IF against Lamin (green) and Vasa (blue) in the larval female germ cells from the indicated species and hybrids and co-stained with DAPI (magenta). Arrows indicate micronuclei. (H) Quantification of micronuclei containing cells in the larval female germ cells from the indicated species and hybrids. **** represents p<0.0001 from Fisher’s exact test. All scale bars are 5μM, yellow dashed lines demarcate nuclear boundary and white dashed lines indicate cell boundary.

The cross between *D. melanogaster* females and *D. simulans*/*D. mauritiana* males yields lethal hybrid males and sterile hybrid females (*12*) (Fig. S1A). Although previous reports have indicated that adult hybrid ovaries are nearly entirely devoid of germ cells (*13, 14*) (Fig. S1B), we found that larval L3 ovaries in hybrids raised at 18°C contained a few surviving Vasa+ germ cells (Fig. S1C, D). We first examined chromocenter formation in female hybrid germ cells using DNA FISH probes against the (AATAT)_n_ repeat (present in both species), the *D. melanogaster*-specific (AATAACATAG)_n_ repeat, and the *D. simulans* species complex-specific (GAACAGAACATGTTC)_n_ repeat.

In the larval ovarian germ cells of pure species, both the *D. melanogaster*-specific (AATAACATAG)_n_ satellite DNA and the *D. simulans* species complex-specific (GAACAGAACATGTTC)_n_ satellite DNA, were clustered into ∼2-3 foci (Fig. 1E, F). In contrast, the larval germ cells of female *D. melanogaster*-*D. simulans* and *D. melanogaster-D. mauritiana* hybrids exhibited increased number of foci, indicating a striking declustering of these satellite DNA repeats (Fig. 1E, F and Fig. S2A). We also observed that hybrid germ cells exhibited declustering of (AATAT)_n_ satellite DNA in comparison to germ cells from pure species (Fig. S2B).

Male hybrids arising from the above cross (Fig. S1A) are larval lethal due to the atrophy of the imaginal discs (*15, 16*) (Table S1). Similar to female hybrid germ cells, we observed defective chromocenter formation in hybrid male imaginal discs in comparison to the pure species controls as indicated by the increased number of satellite DNA foci (Fig. S3A-C, arrows). Interestingly, chromocenter disruption was not observed in the imaginal discs of viable female hybrids: *D. melanogaster*-specific (AATAACATAG)_n_ and *D*.*simulans*-specific (GAACAGAACATGTTC)_n_ satellite DNA were typically associated with each other, revealing intact chromocenter formation, even though these two satellite DNA repeats never co-exist in the cells of pure species (Fig. S3A-C, arrowheads).

Consistent with our previous reports that chromocenter disruption leads to micronuclei formation in *D. melanogaster*, we observed micronuclei in both the hybrid female germ cells(Fig. 1G, H) as well as hybrid male imaginal discs (Fig. S3D, E). In contrast, micronuclei were not observed in the intact imaginal discs of viable female hybrids (Fig. S3D, E). Taken together, our data suggest that cells from atrophied tissues in hybrid animals exhibit defective chromocenter formation and micronuclei. Moreover, these cytological defects are highly correlated with cellular lethality in hybrid animals.

### Removal of hybrid incompatibility genes rescues chromocenter formation

Prior work has identified *D. melanogaster Hmr* (*17, 18*), *D. simulans Lhr* (*19*) and *D. simulans Gfzf* (*20*) as genes causing male lethality in *D*.*melanogaster-D*.*simulans* hybrids. These hybrid incompatibility (HI) genes are considered to act dominantly in hybrids and the removal of even one HI gene restores the viability of male hybrids (*21*). Interestingly, Hmr and Lhr also localize to repetitive DNA and chromocenters in the pure species context (*22*–*25*). Strikingly, we observed significant rescue of chromocenter disruption (Fig. 2A, B, compare to Fig. S3A, B) and micronuclei formation (Fig. 2C, D, compare to Fig. S3D, E) in the imaginal discs of hybrid males, whose viability was restored by mutating *D. simulans Lhr* (Table S1), suggesting that Lhr inhibits chromocenter formation in inviable hybrids. We next used a model of hybrid sterility rescue, where crossing *D. melanogaster In(1)AB, Hmr*^*2*^*/FM6* females to *D. simulans Lhr*^*1*^ males restores ovary development in female hybrids(*26*) (Fig. S4A-C). We found that early germ cells of rescued female hybrids (*In(1)AB, Hmr*^*2*^/+; *Lhr*^*1*^/+) exhibited intact chromocenters (Fig. 2E, F and Fig. S4D, E) and did not form micronuclei (Fig. 2G, H). These results suggested that the incompatibility caused by the HI genes (*Hmr* and *Lhr*) is strongly correlated with chromocenter disruption. In contrast, germ cells of non-rescued sibling controls (*FM6*/+; *Lhr*^*1*^/+) containing Hmr (Fig. S4A-C) exhibited increased declustering of the (AATAT)_n_ satellite DNA (Fig. 2E, F and Fig. S4D, E). Consistent with previous results, chromocenter disruption in the early germ cells of non-rescued hybrid ovaries was accompanied by micronuclei formation (Fig. 2G, H).

**Figure 2.**
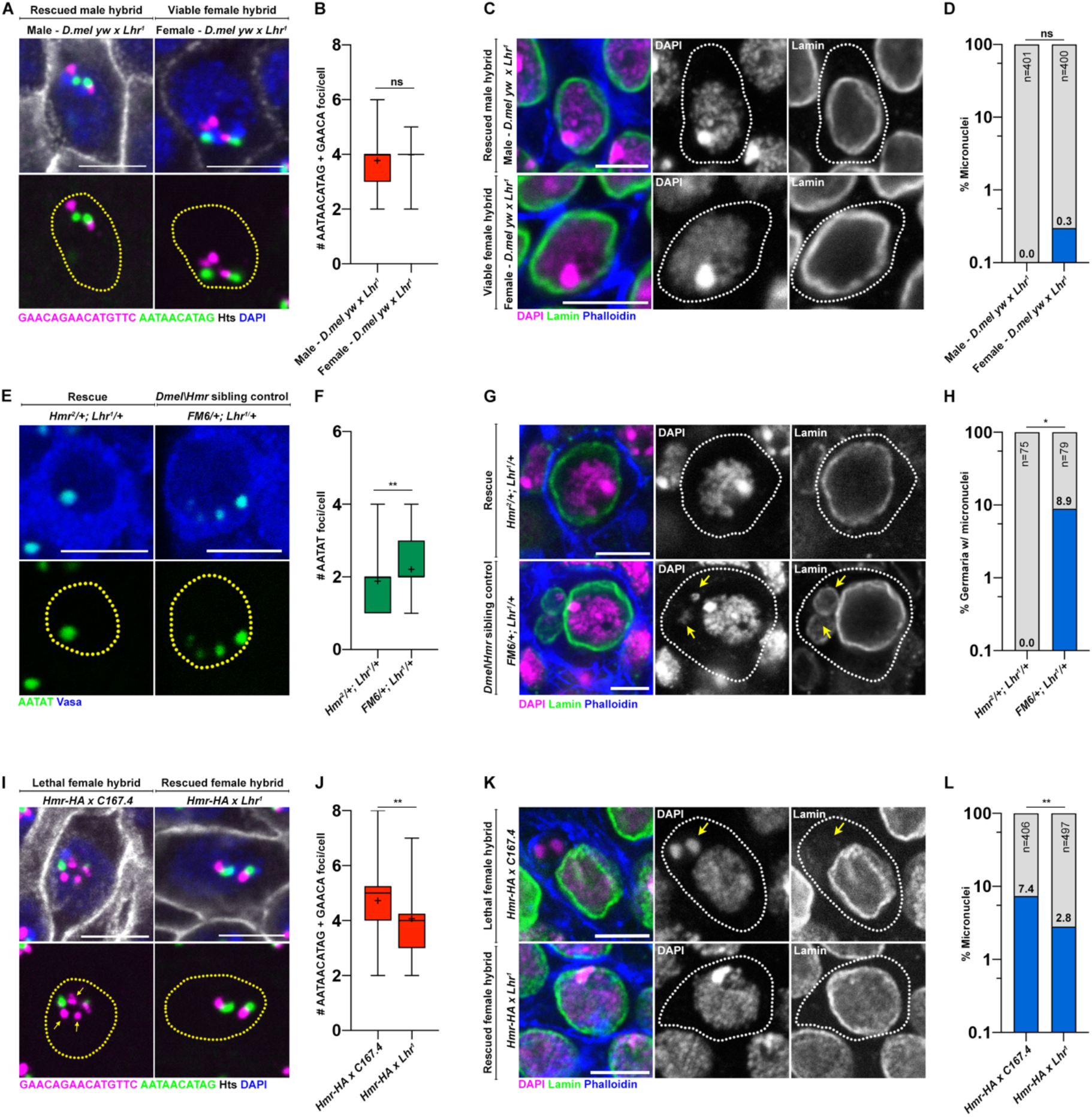
HI factors regulate chromocenter disruption and micronuclei in sterile and lethal hybrids. (A) FISH against the (AATAACATAG)_n_ satellite (green) and the (GAACAGAACATGTTC)_n_ satellite (magenta) on larval imaginal discs from the indicated hybrids and co-stained with DAPI (blue) and Hts (gray). (B) Box-and-whisker plot of total number of (AATAACATAG)_n_ and (GAACAGAACATGTTC)_n_ foci per cell from male (n=63) and female (n=63) *D*.*mel yw x Lhr*^*1*^ hybrids. ns represents p=0.16 from a Student’s t-test and crosshairs mark the mean. (C) IF against Lamin (green) in larval imaginal discs from the indicated hybrids and co-stained with DAPI (magenta) and phalloidin (blue). (D) Quantification of micronuclei containing cells in the larval imaginal discs from male and female *D*.*mel yw x Lhr*^*1*^ hybrids. ns represents p>0.9999 from Fisher’s exact test. (E) FISH against the (AATAT)_n_ satellite (green) in the early germ cells of the indicated 0-2d old adult female hybrids. (F) Box-and-whisker plot of total number of (AATAT)_n_ foci per cell from *In(1)AB, Hmr*^*2*^/+; *Lhr*^*1*^/+ (n=76) and *FM6*/+; *Lhr*^*1*^/+ germ cells (n=73). ** represents p=0.0074 from a Student’s t-test and crosshairs mark the mean. (G) IF against Lamin (green) in early germ cells from the indicated adult 0-2d old female hybrids and co-stained with DAPI (magenta) and phalloidin (blue). Arrows point to micronuclei (H) Quantification of micronuclei containing cells in the early germ cells from the indicated adult 0-2d old female hybrids. * represents p=0.013 from Fisher’s exact test. (I) FISH against the (AATAACATAG)_n_ satellite (green) and the (GAACAGAACATGTTC)_n_ satellite (magenta) on larval imaginal discs from the indicated hybrids and co-stained with DAPI (blue) and Hts (gray). Arrows point to declustered satellite DNA (J) Box- and-whisker plot of total number of (AATAACATAG)_n_ and (GAACAGAACATGTTC)_n_ foci per cell from *Hmr-HA* x *C167*.*4* (n=58) and *Hmr-HA* x *Lhr*^*1*^ (n=62) female hybrids raised at 29°C. ** represents p=0.0022 from a Student’s t-test and crosshairs mark the mean. (K) IF against Lamin (green) in larval imaginal discs from the indicated female hybrids raised at 29°C and co-stained with DAPI (magenta) and phalloidin (blue). Arrows point to micronuclei. (L) Quantification of micronuclei containing cells in larval imaginal discs from the indicate female hybrids. ** represents p=0.0017 from Fisher’s exact test. All scale bars are 5μM, yellow dashed lines demarcate nuclear boundary and white dashed lines indicate cell boundary.

Finally, we found that female hybrids containing an extra copy of *D. melanogaster Hmr* raised at 29°C, which are known to be inviable (*27*) (Table S2), exhibited chromocenter disruption (Fig. 2I, J) and micronuclei formation in the imaginal discs (Fig. 2K, L). Furthermore, these defects were rescued in female hybrids with the *D. simulans Lhr*^*1*^ mutation (Fig. 2I-L, Table S2). Taken together, our data suggest that HI genes trigger chromocenter disruption and micronuclei formation while causing hybrid incompatibility.

### The D1 satellite DNA-binding protein is functionally diverged between species

The above data imply that the inability of chromosomes from two species to form chromocenters may cause cellular defects such as micronuclei, leading to hybrid incompatibility. This raised the question as to whether chromocenter forming proteins such as D1 and Prod, which in *D. melanogaster* bind and cluster the (AATAT)_n_ and (AATAACATAG)_n_ respectively(*5, 6*), might have functionally diverged. Strikingly, we found that the expression of *D*.*simulans* D1 (D1^sim^) failed to rescue the germ cell depletion phenotype of the *D. melanogaster D1* mutant, while expression of *D*.*melanogaster* D1 (D1^mel^) was able to do so (Fig. 3A, B). Moreover, chromocenter formation was clearly impaired in the germ cells of *D*.*melanogaster D1* mutant expressing D1^sim^, in comparison to germ cells expressing D1^mel^ as determined by immunostaining for the D1 and Prod satellite DNA-binding proteins (Fig. 3C-E). Although we did not observe micronuclei in any of the D1^sim^ expressing germ cells, these results reveal a functional divergence between D1^sim^ and D1^mel^ impacting germ cell viability, despite both proteins binding the (AATAT)_n_ satellite DNA (Fig. S5A). Interestingly, we observed that the Prod protein co-localized with clustered (GAACAGAACATGTTC)_n_ satellite DNA in *D. simulans* (Fig. S5B), even though this repeat is highly diverged from the (AATAACATAG)_n_ satellite DNA that is bound by Prod in *D*.*melanogaster*. Surprisingly, *D*.*simulans* Prod (Prod^sim^) was able to fully rescue the loss of viability caused by mutation of *D*.*melanogaster Prod* (Fig. S5C, D) and ectopic expression of Prod^sim^ in *D*.*melanogaster* spermatocytes resulted in chromatin threads connecting heterologous chromosomes, similar to that of Prod^mel^ (*6*) (Fig. S5E), suggesting that Prod from *D*.*simulans* can function in the *D*.*melanogaster* background. We speculate that functional differences of satellite DNA-binding proteins in chromocenter formation may be further exacerbated in hybrids, where the genomes of both species are brought together in the same nucleus.

**Figure 3.**
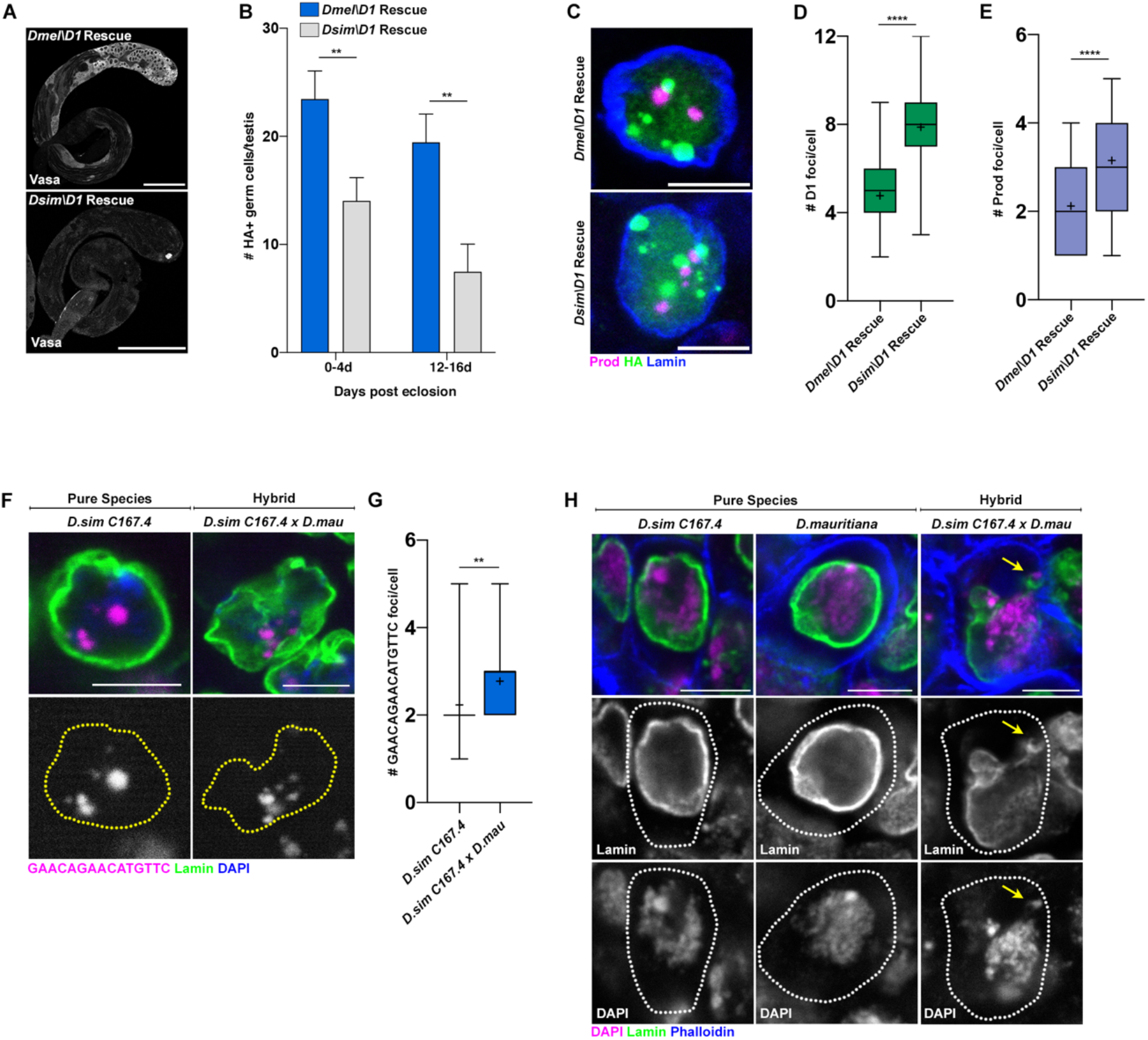
Functional divergence of the D1 chromocenter forming protein from *D*.*melanogaster* and *D*.*simulans*. *D1* mutant (*D1*^*LL03310*^) testes expressing HA-tagged *UASt-D1*^*mel*^ and *UASt-D1*^*sim*^ under the control of *nos-gal4* were stained for Vasa (gray). Scale bars are 25μM. (B) Quantification of HA+ germ cells in testes from 0-4d old and 12-16d old males of the following genotypes, *nos>D1*^*mel*^; *D1*^*LL03310*^ (n=29, 0-4d) and (n=31, 12-16d) and *nos>D1*^*sim*^; *D1*^*LL03310*^ (n=31, 0-4d) and n=32, 12-16d). ** represents p=0.006 (0-4d) and p=0.001 (12-16d) from a Student’s t-test. (C) IF against Prod (magenta), HA (green) and Lamin (blue) in spermatogonial cells of the indicated genotypes. (D) Box-and-whisker plot of HA foci/cell in *D1* mutant (*D1*^*LL03310*^) spermatogonia expressing *D1*^*mel*^*-HA* (n=57) or *D1*^*sim*^*-HA* (n=60). **** represents p<0.0001 from Student’s t-test and crosshairs mark the mean. (E) Box-and-whisker plot of Prod foci/cell in *D1* mutant (*D1*^*LL03310*^) spermatogonia expressing *D1*^*mel*^*- HA* (n=57) or *D1*^*sim*^*-HA* (n=60). **** represents p<0.0001 from Student’s t-test and crosshairs mark the mean. (F) FISH against the (GAACAGAACATGTTC)_n_ satellite (magenta) in spermatogonial cells from the indicated 0-3d old males and co-stained with DAPI (blue) and Lamin (green). (G) Box-and-whisker plot of large autosomal (GAACAGAACATGTTC)_n_ foci per spermatogonial cell from *D*.*sim C167*.*4* (n=39) and *D*.*sim C167*.*4* x *D*.*mau w*^*+*^ (n=31). ** represents p=0.0067 from a Student’s t-test and crosshairs mark the mean. (H) IF against Lamin (green) in spermatogonial from the indicated 0-3d old males and co-stained with DAPI (magenta) and phalloidin (blue). Arrows point to micronuclei. All scale bars (except panel A) are 5μM and yellow dashed lines demarcate nuclear boundary.

### Chromocenter disruption is a common phenotype among *Drosophila* hybrids

Due to their recent divergence ∼250,000 years ago, the satellite DNA content of *D. simulans* and *D. mauritiana* are more similar to each other in comparison to *D. melanogaster* (*10*). However, a few satellite DNA remain distinct between these species (e.g. the *D. simulans* contains Y-specific (AATAAAC)_n_ and (AAGAGAG)_n_ repeats, which are lacking in *D. mauritiana*) while other satellite DNA show different abundances and locations on the chromosomes e.g. the (GAACAGAACATGTTC)_n_ and (AATAG)_n_ repeats (*10*). Crosses between *D. simulans* females and *D. mauritiana* males results in fertile female progeny but sterile male progeny (Fig. S6A), with the testes of sterile males reported to exhibit loss of premeiotic germ cells (*28, 29*). We confirmed that the testes of male hybrids exhibited dramatic early germ cell loss with age (Fig. S6B). Moreover, we observed chromocenter disruption (Fig. 3F, G) and micronuclei formation (Fig. 3H), specifically in the male germ cells of hybrid testes (0% MN in *D*.*sim C167*.*4* (n=18) vs. 38.5% in *D*.*sim C167*.*4* x *D*.*mau w*^*+*^ hybrids (n=26), p=0.0027 from Fisher’s exact test.) In contrast, germ cell content and tissue morphology were intact in the fertile female hybrids between *D. simulans* and *D. mauritiana* (Fig. S6C). Therefore, chromocenter disruption due to differences in satellite DNA composition, even between very recently diverged species, may drive micronuclei formation and cell death and promote hybrid incompatibility.

The divergence of satellite DNA repeats between species has been long postulated to mediate hybrid incompatibility (HI) and reproductive isolation. However, poor conservation of these non-coding repeats across species is also one of the main reasons why satellite DNA is typically considered to be ‘junk DNA’, making it difficult to speculate on the ‘incompatibility of (useless) junk’. Our recent work identified a conceptual and mechanistic framework to understand how satellite DNA functions within species: pericentromeric satellite DNA association between heterologous chromosomes (chromocenters) bundles the entire set of chromosomes within the nucleus(*5, 6*). In this study, we demonstrate that this framework can explain how divergent satellite DNA repeats impede the viability and fertility of hybrids. Indeed, we provide the first proof-of-concept evidence that cells from hybrid animals containing distinct satellite DNA repeats exhibit phenotypes unique to chromocenter disruption. We further suggest that differential evolutionary trajectories of chromocenter-associated proteins in response to the rapid turnover of satellite DNA repeats within species can plant the seed for subsequent reproductive isolation and speciation. Together, our study lays the foundation for understanding hybrid incompatibility at a cellular level in Drosophila as well as other eukaryotes.

## Acknowledgements

We thank Dan Barbash, Trisha Wittkopp, Tibor Torok, the Bloomington Drosophila Stock Center, the Kyoto Stock Center, the Drosophila Species Stock Center and the Developmental Studies Hybridoma Bank for reagents and resources. We thank members of the Yamashita lab and Jagannathan lab for discussion and comments on the manuscript. This research was supported by the Howard Hughes Medical Institute (YY) and an American Heart Association postdoctoral fellowship (MJ). MJ and YY conceived the project, interpreted the data and wrote the manuscript. Both authors contributed to conducting experiments and analyzing data.

