## Supplementary Materials for "Defective satellite DNA clustering into chromocenters underlies hybrid incompatibility in *Drosophila*"

### Materials and Methods

**Fly husbandry and strains.** All fly stocks were raised on standard Bloomington medium at 25°C unless otherwise indicated. *D. melanogaster yw* was used as a wild type stock. The following fly stocks were obtained from the Drosophila species stock center: *D. simulans* $w^{501}$ (DSSC#14021-0251.195), *D. mauritiana* $w^{1}$ (DSSC#14021-0241.60), *D.mauritiana* $w^{+}$ wild type (DSSC#14021-0241.150), *D. simulans* $Lhr^{1}$ (DSSC# 14021-0251.023). $D1^{LL03310}$ (DGRC140754), *FRT42D* $prod^{k08810}$ (DGRC111248) and D.simulans C167.4 (DGRC107850) were obtained from the Kyoto stock center. $prod^{U}$ (BDSC42686) was obtained from the Bloomington *Drosophila* stock center. *Hmr-HA* (*25*) and *In(1)AB, Hmr2/FM6* (*26*) were gifts from Daniel Barbash while the *D.simulans Tsimbazaza* strain (*13*) was a gift from Patricia Wittkopp. *nos-gal4* (*30*) and *bam-gal4* (*31*) and *pUASt-GFP-*$Prod^{mel}$ (*6*) have been previously described.

**Transgene construction.** For construction of *pUASt-GFP-*$Prod^{sim}$, a codon optimized $Prod^{sim}$ ORF was subcloned into the NotI and KpnI sites of *pUASt-EGFP-attB* (*32*) resulting in *pUASt-GFP-*$Prod^{sim}$. For construction of *p400-GFP-*$Prod^{sim}$, a 400bp promoter upstream of the Prod start site from *D.melanogaster* was PCR amplified using the following primer pair, GATCAAGCTTCTGTTGTTATGCATATCGTTC and GATCGAATTCCCGGGTATCCTTGCTC and subcloned into the HindIII and EcoRI sites on *pUASt-GFP-*$Prod^{sim}$, replacing the UAS sequence. Transgenic flies were generated for both plasmids using PhiC31 integrase-mediated transgenesis into the *attP40* site (BestGene). The following plasmids, *pUASt-D1mel-HA* and *pUASt-D1sim-HA* were obtained from Dan Barbash (*33*) and transgenic flies were generated using PhiC31 integrase-mediated transgenesis into the *attP2* site (BestGene).

**Immunofluorescence staining and microscopy.** For *Drosophila* tissues, immunofluorescence staining was performed as described previously (*34*). Briefly, tissues were dissected in PBS, transferred to 4% formaldehyde in PBS and fixed for 30 minutes. Tissues were then washed in PBS-T (PBS containing 0.1% Triton-X) for at least 60 minutes, followed by incubation with primary antibody in 3% bovine serum albumin (BSA) in PBS-T at 4°C overnight. Samples were washed for 60 minutes (three 20-minute washes) in PBS-T, incubated with secondary antibody in 3% BSA in PBS-T at 4°C overnight, washed as above, and mounted in VECTASHIELD with DAPI (Vector Labs). The following primary antibodies were used: rabbit anti-vasa (1:200; d-26; Santa Cruz Biotechnology), rat anti-Vasa (Developmental Studies Hybridoma Bank), mouse anti-LaminDm$_0$ (ADL84.12, 1:200, Developmental Studies Hybridoma Bank), mouse anti-Hts (1B1, Developmental

Studies Hybridoma Bank), rat anti-HA (Sigma, 3F10 ), Phalloidin-Alexa488 (Abcam, ab176753 ), Phalloidin-Alexa546 (ThermoFisher, a22283, 1:200), rabbit anti-Prod (gift from Tibor Torok, 1:5000) and guinea pig anti-D1 (generated using the synthetic peptide CDGENDANDGYVSDNYNDSESVAA (Covance)). Fluoresecent images were taken using a Leica TCS SP8 confocal microscope with 63x oil-immersion objectives (NA=1.4). Brightfield images were acquired using a Keyence microscope. Images were processed using Adobe Photoshop software.

**DNA fluorescence *in situ* hybridization.** Whole mount *Drosophila* tissues were prepared as described above, and optional immunofluorescence staining protocol was carried out first. Subsequently, samples were post-fixed with 4% formaldehyde for 10 minutes and washed in PBS-T for 30 minutes. Fixed samples were incubated with 2 mg/ml RNase A solution at 37°C for 10 minutes, then washed with PBS-T + 1mM EDTA. FISH using heat denaturation was carried out as follows: samples were washed in 2xSSC-T (2xSSC containing 0.1% Tween-20) with increasing formamide concentrations (20%, 40% and 50%) for 15 minutes each followed by a final 30-minute wash in 50% formamide. Hybridization buffer (50% formamide, 10% dextran sulfate, 2x SSC, 1mM EDTA, 1 μM probe) was added to washed samples. Samples were denatured at 91°C for 2 minutes, then incubated overnight at RT. For hybrid tissues, FISH using acid denaturation was used instead of heat denaturation and carried out as follows: samples were washed in 2xSSC-T, DNA was denatured using a 30-minute incubation with 2N HCl at RT followed by three rinses in ice-cold PBS-T. Hybridization buffer (60% formamide, 10% dextran sulfate, 2x SSC, 1mM EDTA, 0.5-1μM probe) was added to washed samples and incubated overnight at RT. For mitotic chromosome spreads, larval 3$^{rd}$ instar brains were squashed according to previously described methods (*35*). Briefly, tissue was dissected into 0.5% sodium citrate for 5-10 minutes and fixed in 45% acetic acid/2.2% formaldehyde for 4-5 minutes. Fixed tissues were firmly squashed with a cover slip and slides were submerged in liquid nitrogen until bubbling ceased. Coverslips were then removed with a razor blade and slides were dehydrated in 100% ethanol for at least 5 minutes. After drying, hybridization mix (50% formamide, 2x SSC, 10% dextran sulfate, 100 ng of each probe) was applied directly to the slide, samples were heat denatured at 95°C for 5 minutes and allowed to hybridize overnight at room temperature. Following hybridization, slides were washed 3 times for 15 minutes in 0.2X SSC and mounted with VECTASHIELD with DAPI (Vector Labs). The following probes were used for *Drosophila* in situ hybridization: (AATAT)$_6$, (AATAACATAG)$_3$ and (GAACAGAACATGTTCGAACAGAACATGTTCGAACA) and have been previously described (*10*).

**Rescue experiments.** Rescue with $D1^{sim}$: The *pUASt-D1mel-HA* and *pUASt-D1sim-HA* transgenes were each recombined with the $D1^{LL03310}$ mutant allele and crossed to *nos-gal4::VP16*; $D1^{LL03310}$ flies at room temperature. Testes from *nos>UAS-*$D1^{mel}$; $D1^{LL03310}$ and *nos>UAS-D1sim*; $D1^{LL03310}$ adult males were dissected and the number of HA+ germ cells were scored to assess the extent of rescue. The $Prod^{sim}$ rescue allele was generated by recombining the *p400-GFP-*$Prod^{sim}$ transgene with $prod^{k08810}$. The $Prod^{sim}$ rescue allele and the *FRT42D* $prod^{k08810}$ allele as a control were each crossed to $prod^{U}$ in vials at 25°C. The percent of trans-heterozygous *prod* mutant flies was quantified in each replicate.

**Table S1. Viability of male and female hybrids in crosses between *D.melanogaster* females and *D.simulans/D.mauritiana* males.** Approximately 20 males and females were crossed with each other in bottles and the number of progeny was quantified from the indicated inter-specific crosses at 18ºC and 25ºC. Numbers in parentheses indicate the number of crosses performed.

| Cross | # Male Progeny | % Male Progeny | # Female Progeny | % Female Progeny | Total Progeny |
|---|---|---|---|---|---|
| *yw* 25ºC (4) | 750 | 55.9 | 591 | 44.1 | 1341 |
| *yw* 18ºC (2) | 269 | 47.4 | 299 | 52.6 | 568 |
| *yw* x *D.simulans C167.4* 25 ºC (5) | 0 | 0.0 | 684 | 100.0 | 684 |
| *yw* x *D.simulans C167.4* 18 ºC (4) | 0 | 0.0 | 514 | 100.0 | 514 |
| *yw* x *D.mauritiana* 25 ºC (7) | 1 | 0.1 | 1275 | 99.9 | 1276 |
| *yw* x *D.mauritiana* 18 ºC (2) | 0 | 0.0 | 452 | 100.0 | 452 |
| *yw* x *D.simulans Lhr*$^{1}$ 25 ºC (4) | 405 | 47.2 | 453 | 52.8 | 858 |
| *yw* x *D.simulans Lhr*$^{1}$ 18 ºC (2) | 101 | 34.4 | 193 | 65.6 | 294 |
| *yw* x *D.simulans Tsimbazaza* 18 ºC (3) | 0 | 0.0 | 370 | 100.0 | 370 |

**Table S2. Female lethality in hybrids carrying an extra copy of *D.melanogaster Hmr*.** Approximately 20 males and females were crossed with each other in bottles and the number of progeny was quantified from the indicated inter-specific crosses at 29ºC. Numbers in parentheses indicate the number of crosses performed.

| Cross | # Male Progeny | # Female Progeny | Total Progeny |
|---|---|---|---|
| *D.mel Hmr-HA* x *D.sim C167.4* (2) | 0 | 80 | 80 |
| *D.mel Hmr-HA* x *D.sim* $Lhr^{1}$ (2) | 0 | 4 | 4 |

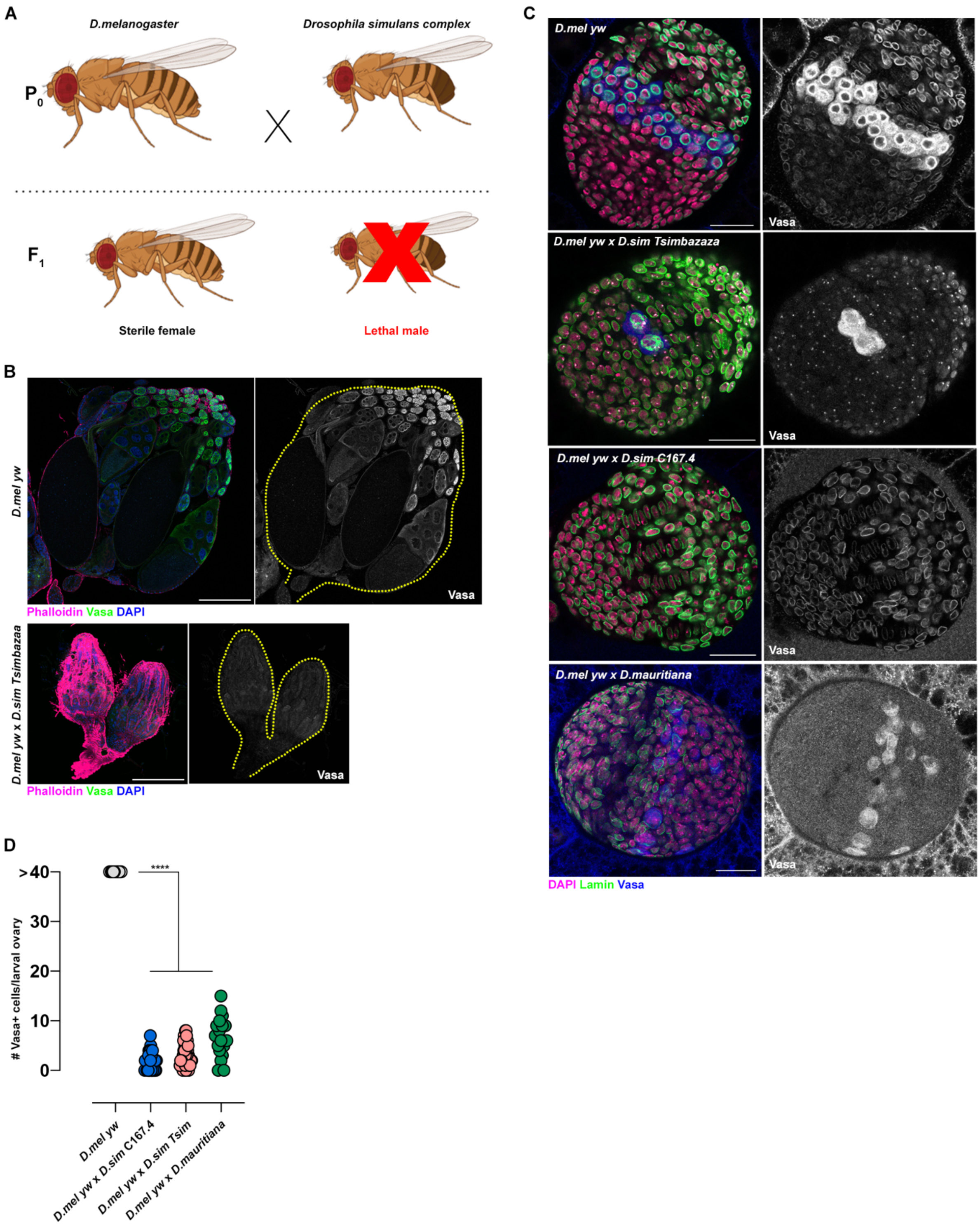


**Figure S1. Surviving germ cells in female larval hybrids raised at 18°C.** (A) Crosses between *D.melanogaster* females and males from the *D.simulans* complex males yields sterile female progeny and inviable male progeny. (B) IF against Vasa (green) in adult ovaries from the indicated species and hybrids raised at RT and co-stained with DAPI (blue) and phalloidin (magenta). Scale bars are 100µM (C) IF against Vasa (blue) and Lamin (green) in larval ovaries from the indicated species and hybrids raised at 18°C and co-stained with DAPI (magenta). Scale bars are 25µM. (D) Quantification of

Vasa+ germ cells per larval ovary from *D.mel yw* (n=13), *D.mel yw* x *C167.4* hybrid females (n=39), *D.mel yw* x *D.sim Tsimbazaza* hybrid females (n=33) and *D.mel yw* x *mau* hybrid females (n=27). **** represents $p<0.0001$ based on Tukey's multiple comparisons test from an ordinary one-way ANOVA.

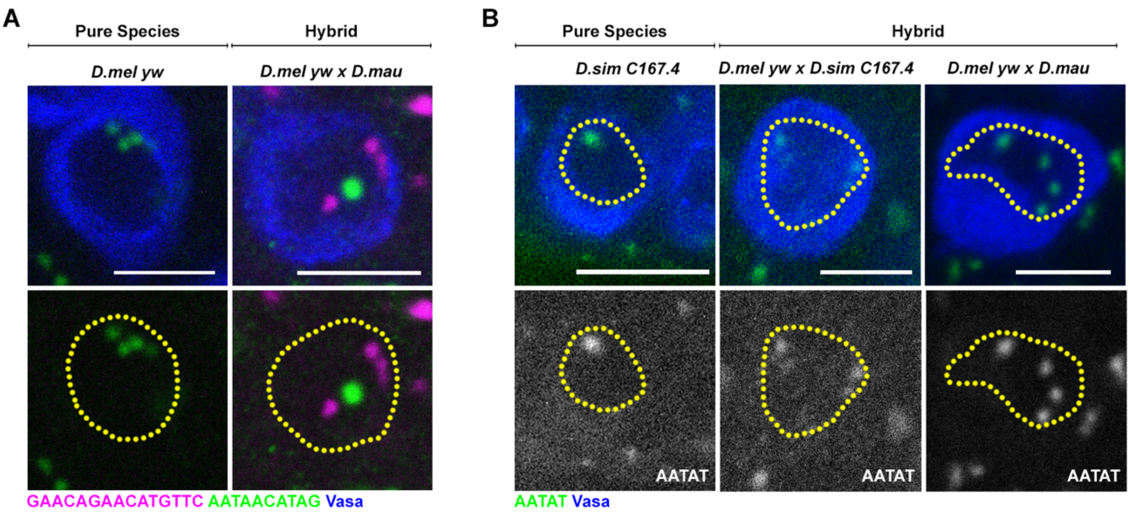


**Figure S2. Chromocenter disruption in the larval germ cells of *D.melanogaster-D.simulans/D.mauritiana* hybrids.** (A) FISH against the $(AATAACATAG)_n$ satellite (green) and the $(GAACAGAACATGTTC)_n$ satellite (magenta) in the larval female germ cells from the indicated species and hybrids and co-stained with Vasa (blue). (B) FISH against the $(AATAT)_n$ satellite (green) in the larval female germ cells from the indicated species and hybrids and co-stained with Vasa (blue). All scale bars are 5μM and yellow dashed lines demarcate nuclear boundary.

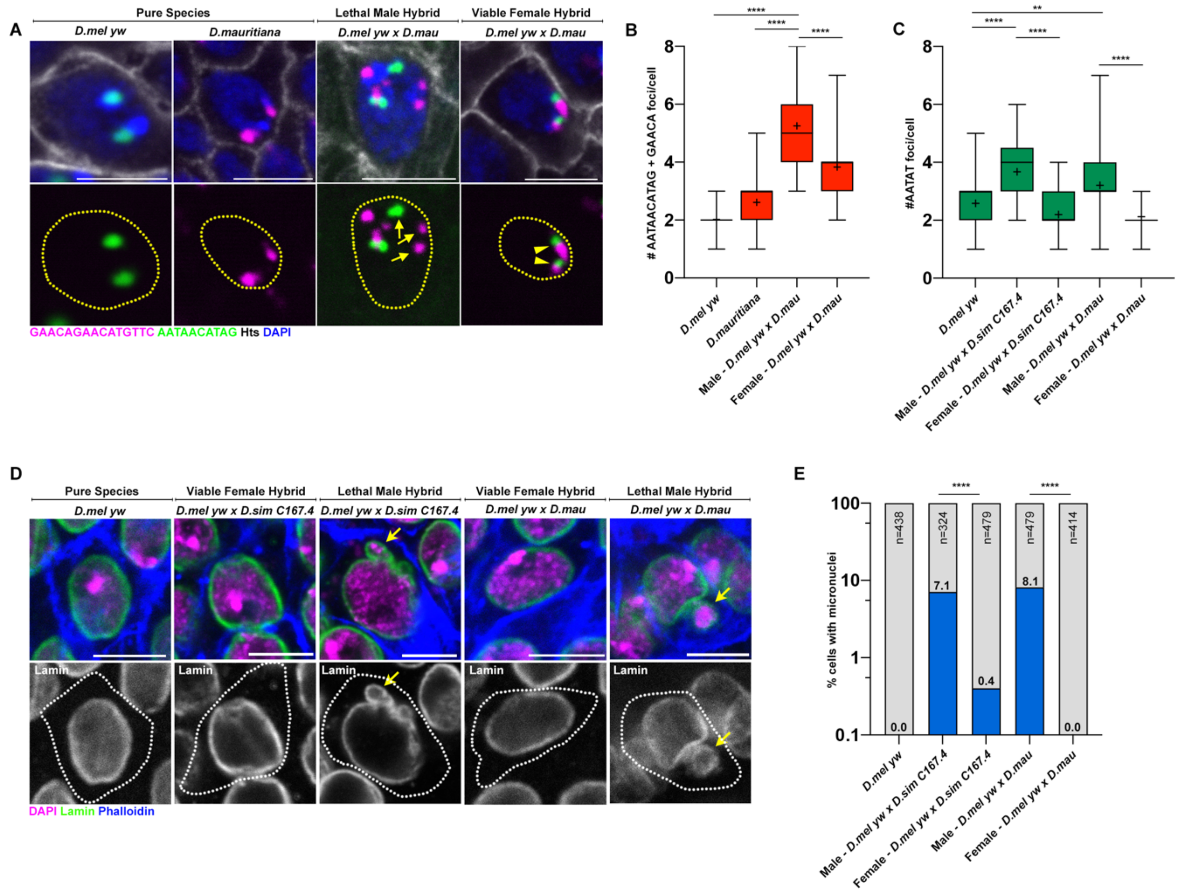


**Figure S3. Chromocenter disruption and micronuclei in the larval imaginal discs of lethal male *D.melanogaster-D.simulans/D.mauritiana* hybrids.** (A) FISH against the $(AATAACATAG)_n$ satellite (green) and the $(GAACAGAACATGTTC)_n$ satellite (magenta) on larval imaginal discs from the indicated species and hybrids and co-stained with DAPI (blue) and Hts (gray). (B) Box-and-whisker plot of total number of $(AATAACATAG)_n$ and $(GAACAGAACATGTTC)_n$ foci per cell from *D.mel yw* (n=46), *D.mau* (n=64) and male (n=43) and female (n=60) *D.mel yw* x *D.mau* hybrids. **** represents p<0.0001 based on Tukey's multiple comparisons test from an ordinary one-way ANOVA and crosshairs mark the mean. (C) Box-and-whisker plot of total number of $(AATAT)_n$ foci per cell from *D.mel yw* (n=48), male (n=37) and female (n=51) *D.mel yw* x *D.sim C167.4* hybrids and male (n=39) and female (n=41) *D.mel yw* x *D.mau* hybrids. ** represents p=0.0049 and **** represents p<0.0001 based on Tukey's multiple comparisons test from an ordinary one-way ANOVA and crosshairs mark the mean. (D) IF against Lamin (green) in larval imaginal discs from the indicated species and hybrids and co-stained with DAPI (magenta) and phalloidin (blue). (E) Quantification of micronuclei containing cells in the larval imaginal discs from the indicated species and hybrids. **** represents p<0.0001 from Fisher's exact test. All scale bars are 5μM, yellow dashed lines demarcate nuclear boundary and white dashed lines indicate cell boundary.

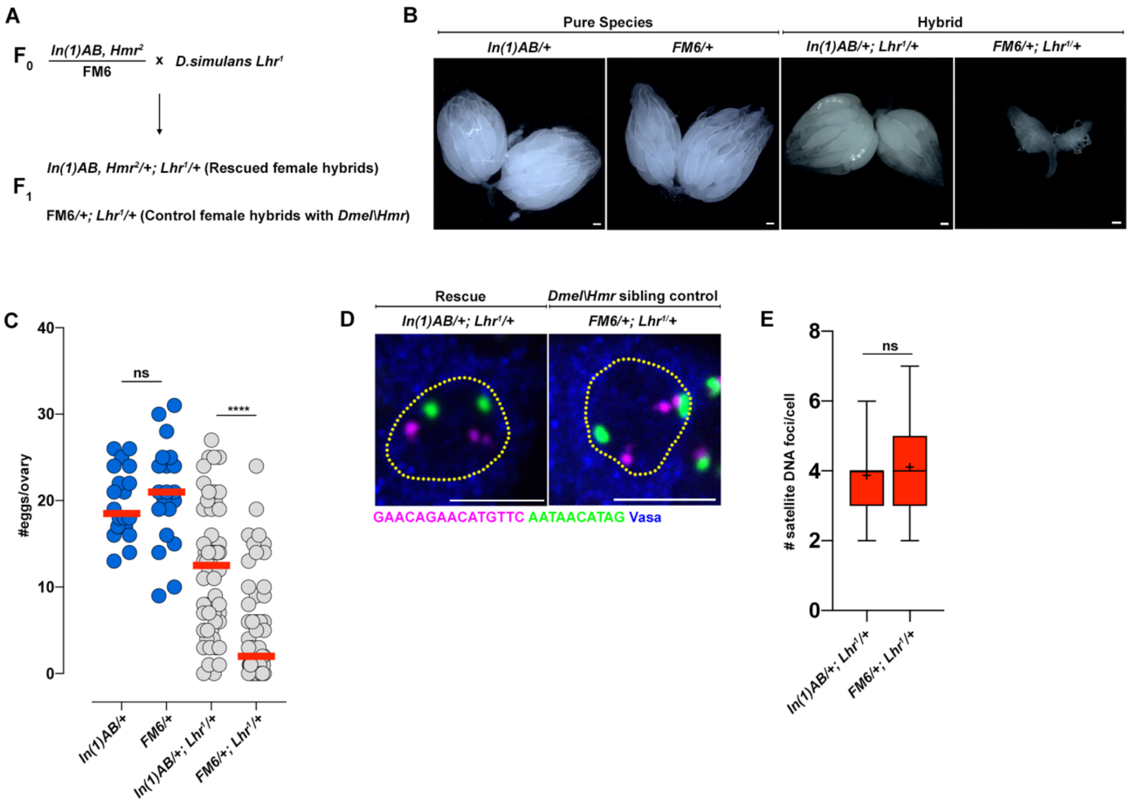


**Figure S4. Rescue of hybrid female germ cell development using *In(1)AB, Hmr*$^{2}$ and *Lhr*$^{1}$.** (A) Crossing scheme for hybrid germ cell rescue. (B) Brightfield images of ovaries from 3-7d old females of pure species, *In(1)AB, Hmr*$^{2}$/+ and *FM6*/+ (obtained by crossing *In(1)AB, Hmr*$^{2}$*/FM6* x *D.mel yw*) and of hybrids, *In(1)AB, Hmr*$^{2}$/+; *Lhr*$^{1}$/+ and *FM6*/+; *Lhr*$^{1}$/+ (obtained by crossing *In(1)AB, Hmr*$^{2}$*/FM6* x *D.sim Lhr*$^{1}$*).* Scale bar is 100µM. (C) Quantification of eggs/ovary from the following genotypes, *In(1)AB, Hmr*$^{2}$/+ (pure species, n=20) and *FM6*/+ (pure species, n=20) and *In(1)AB, Hmr*$^{2}$/+; *Lhr*$^{1}$/+ (hybrid, n=56) and *FM6*/+; *Lhr*$^{1}$/+ (hybrid, n=54). ns represents p=0.9455 and **** represents p<0.0001 based on Tukey's multiple comparisons test from an ordinary one-way ANOVA and the red lines mark the median. (D) FISH against the $(AATAACATAG)_n$ (green) and $(GAACAGAACATGTTC)_n$ satellite (magenta) in the early germ cells of the indicated 0-2d old adult female hybrids co-stained with Vasa (blue). (E) Box-and-whisker plot of total number of $(AATAACATAG)_n$ and $(GAACAGAACATGTTC)_n$ foci per cell from *In(1)AB, Hmr*$^{2}$/+; *Lhr*$^{1}$/+ (n=47) and *FM6*/+; *Lhr*$^{1}$/+ germ cells (n=54). ns represents p=0.2284 from a Student's t-test and crosshairs mark the mean.

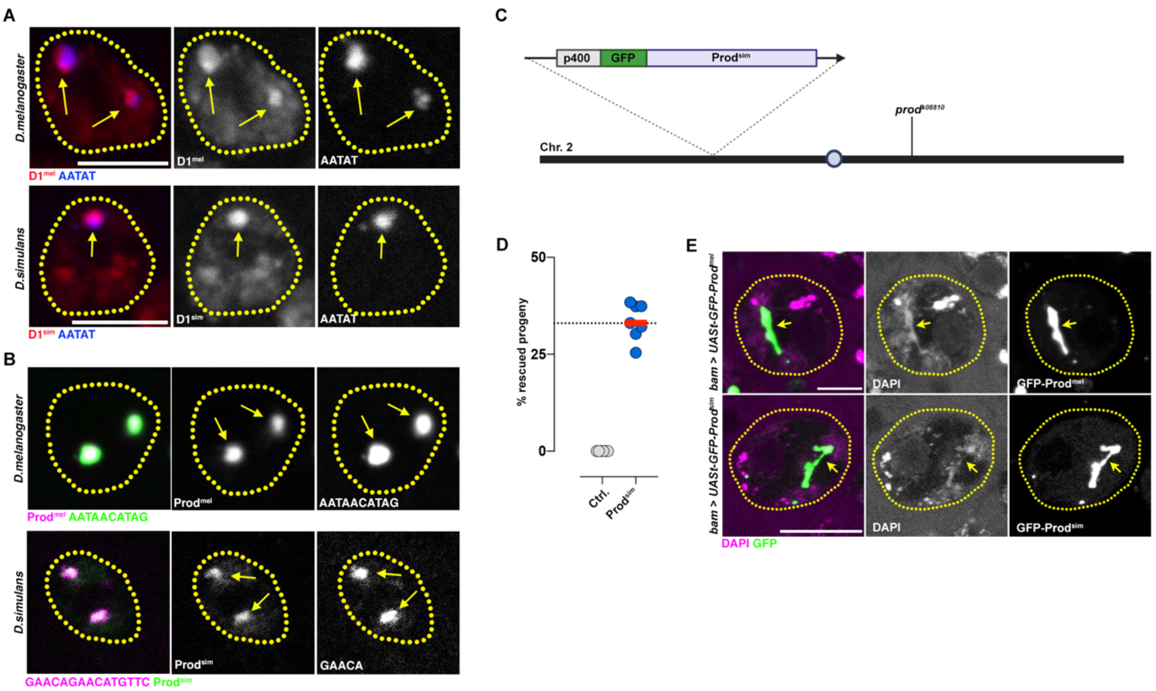


**Figure S5. Expression and functional complementation of the D1 and Prod satellite DNA-binding proteins from *D.melanogaster* and *D.simulans*.** (A) FISH against the $(AATAT)_n$ satellite (blue) in spermatogonial cells of *D.mel yw* and *D.sim Tsimbazaza* testes co-stained for D1(red). (B) FISH against the $(AATAACATAG)_n$ satellite (green) and $(GAACAGAACATGTTC)_n$ satellite (magenta) in spermatogonial cells of *D.mel yw* and *D.sim Tsimbazaza* testes co-stained for Prod (magenta/green). (C) Schematic of p400-GFP-$Prod^{sim}$ rescue allele: A 400bp promoter sequence upstream of the $Prod^{mel}$ start site was placed upstream of a GFP-tagged and codon optimized $Prod^{sim}$ sequence, inserted into the attP40 locus and recombined with the *prod*$^{k08810}$ loss-of-function allele. (D) Rescue of prod mutant (*prod*$^{k08810/U}$) lethality was assessed using p400-GFP-$Prod^{sim}$, *prod*$^{k08810}$ (Rescue allele, 757 flies were counted from 7 crosses) and *prod*$^{k08810}$ (Ctrl, 230 flies were counted from 6 crosses). Red line indicates median while dashed line indicates percent progeny for complete rescue. (E) Ectopic expression of *UASt-GFP-Prod*$^{mel}$ and *UASt-GFP-Prod*$^{sim}$ in *D.melanogaster* spermatocytes under the control of *bam-gal4* co-stained with DAPI. Arrows indicate Prod+ threads connecting autosomal chromosome territories. All scale bars are 5µM and yellow dashed lines demarcate nuclear boundary.

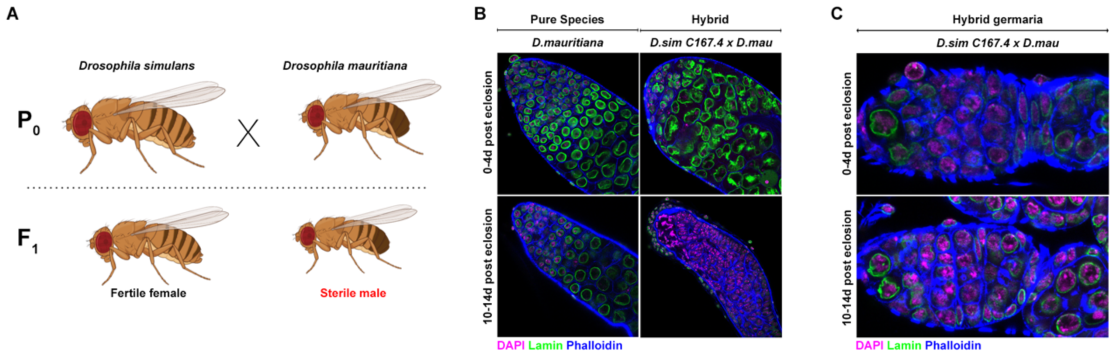


**Figure S6. Testes from D.simulans-D.mauritiana hybrids exhibit rapid germ cell loss.** (A) Crosses between *D.simulans* females and *D.mauritiana* males yield sterile male progeny and fertile female progeny. (B) IF against Lamin (green) in testes from 0-4d old and 10-14d old adults from the indicated species and hybrids raised at 25°C and co-stained with DAPI (magenta) and phalloidin (blue). (C) IF against Lamin (green) in germaria from 0-4d old and 10-14d old adult females from the indicated species and hybrids raised at 25°C and co-stained with DAPI (magenta) and phalloidin (blue).
